# The six steps of the F_1_-ATPase rotary catalytic cycle

**DOI:** 10.1101/2020.12.15.422962

**Authors:** Meghna Sobti, Hiroshi Ueno, Hiroyuki Noji, Alastair G. Stewart

## Abstract

F_1_F_o_ ATP synthase interchanges phosphate transfer energy and proton motive force via a rotary catalysis mechanism. When isolated, its F_1_-ATPase catalytic core can hydrolyze ATP, rotating its γ rotor subunit. Although previous structural studies have contributed greatly to understanding rotary catalysis in F_1_, the structure of one major conformational state detected in single-molecule studies, termed the binding dwell state, has not yet been determined. Here, by exploiting a temperature-sensitive F_1_-ATPase mutant from *Bacillus* PS3, the structure of this binding dwell state was established together with that of the catalytic dwell state. Each state showed three catalytic β subunits in different conformations, providing the complete set of six β subunit conformational states taken up during catalysis cycle. These structures provide molecular details for the power-stroke conformational change that occurs upon ATP binding and induces a ~80° γ subunit rotation, as well as a second torque-generating conformational change, triggered by hydrolysis and product release, that produces a ~40° rotation. This study also identifies a putative phosphate-releasing tunnel that indicates how ADP and phosphate releasing steps are coordinated. Overall these findings provide a structural basis for the entire F_1_-ATPase rotary catalysis cycle.

## Main

F_1_F_o_ ATP synthase, a key component in the generation of cellular metabolic energy, is a biological rotary motor that converts proton motive force (pmf) to adenosine tri-phosphate (ATP) in both oxidative phosphorylation and photophosphorylation^1–4^. The enzyme is comprised of two rotary motors, the F_1_-ATPase and F_o_ motor, that are coupled together by two stalks: a central γ subunit “rotor” stalk and a peripheral “stator” stalk. The F_o_ motor spans the membrane and converts the potential energy from the pmf into mechanical rotation of the central rotor. This central rotor drives conformational changes in the catalytic F_1_-ATPase that generate ATP from ADP and inorganic phosphate (P_i_)^5,6^. Studies of the isolated F_1_-ATPase have shown it can also function as a molecular motor, driven by ATP hydrolysis^3,5,7^. The F_1_-ATPase consists of three α and three β subunits, arranged in an alternating ring. The three catalytic sites are at the interface between the α and β subunits, with the β subunit being the principal catalytic subunit that undergoes conformational changes to drive the rotation of the γ subunit^5–9^. The pseudo 3-fold architecture of the enzyme dictates that the γ subunit rotates 120° for each ATP turned over.

Single molecule studies on the F_1_-ATPase initially showed a principal unitary γ subunit rotation step of 120° that is coupled with a single turn-over of ATP hydrolysis^10^. This 120° step was further resolved into two discrete substeps of ~80° and ~40°^11–13^, although the *Paracoccous* enzyme does not show an obvious substep^14^. The rotation dwells before the 80° and 40° substeps are termed the “binding dwell” and “catalytic dwell”, respectively, because these dwells were first identified as either waiting for ATP binding^11^ or hydrolysis^15,16^. Subsequently, rotation assays showed that phosphate is likely released during the 40° substep^17^ and experiments using fluorescent nucleotide showed that ADP is released during the 80° step^17,18^. All these findings can be summarized into a circular reaction scheme that shows the major catalytic events linked to rotary position (Fig. 1). In a full rotation, F_1_ makes conformational transitions between the binding dwell and catalytic dwell states, with each β subunit undergoing six conformational transitions.

**Figure 1:**
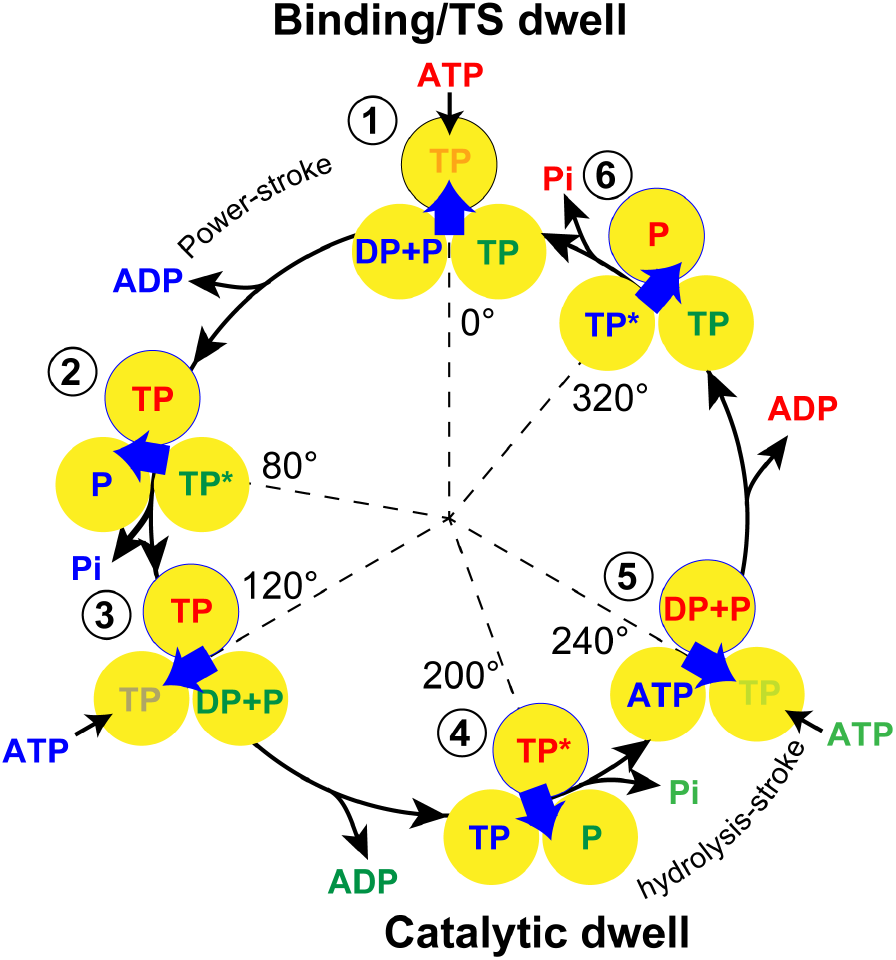
Rotation reaction scheme of F_1_-ATPase suggested by single molecule experiments. β subunits are depicted as yellow circles with binding site occupancy labelled, the γsubunit is depicted as blue arrow and, for clarity, the α subunits are not shown. The F_1_-ATPase rotates with two sub-strokes each of 80 °and 40 °, respectively. The ATP binding dwell precedes ATP binding and is followed by the “power-stroke”, whereas the catalytic dwell precedes the “hydrolysis-stroke. Single β subunits are outlined in black and numbered from 1 to 6 at each dwell to highlight the reaction path.

Bovine mitochondrial F_1_-ATPase (bMF_1_) has been a long-standing model system to characterize the F_1_-ATPase structurally. Although rich in biochemical information^5,19–21^, the structures of bMF_1_ principally represent the catalytic dwell state, or related transition states, with the position of the γ subunit determined by crystal lattice contacts with the crown region of adjacent bMF_1_ complexes^20^. In the ground state structure of bMF_1_^19^, each β subunit contained a different nucleotide composition and was either in an “open” or “closed” conformation, with the subunits named due to the nucleotide occupancy: β_DP_ contained MgADP and was in a closed state, β_TP_ contained MgAMP-PNP and was in a closed state and β_E_ was in an open state with no nucleotide bound. The β_DP_ and β_TP_ subunits had essentially the same conformation, though the αβ interface is more closed in the αDP-β_DP_ pair. The β_E_ subunit assumes the open conformation, hinging its C-terminal domain outward. Overall, the crystal structures of bMF_1_ in the ground state represent the catalytic dwell state and show three of the six conformational states of the β subunit found in the single molecule rotation assays.

Structural studies that attempt to resolve structures other than the ground state have also been reported. In the structure of bMF_1_ solved in the presence of aluminum fluoride, the β_E_ subunit binds to ADP and sulfate, adopting a “half-closed” conformation, *i.e.* in an intermediate conformation between the closed and open conformations^22^. The γ subunit was rotated by −15° from the rotary position seen in the ground state^23^. Based on these features, this structure is considered to represent a pre-ADP intermediate state of F_1_ that appears in the 80° substep, *i.e.* in the conformational transition from the binding dwell state to the catalytic dwell state. Autoinhibited structures of *E. coli* F_1_-ATPases (EF_1_) show the β subunits in half-closed, closed, open conformations^24,25^, and the autoinhibited structures of *Bacillus* PS3 F_1_-ATPase (TF_1_) show the β subunits in open, closed, open conformations^26,27^, with all of these studies showing the inhibitory ε subunit in an “up” position that is believed to prevent rotation. Structures of yeast^28^ and bacterial^29^ F_1_-ATPases also show a similar rotational state to the bMF_1_ ground state. To date, no structure representing the binding dwell has been obtained. Thus, the structures of three of the six conformational states of the β subunit during ATP catalysis have not been established, and it remains unclear how the β subunit executes power-strokes upon ATP binding or catalysis and product release.

The higher stability of the catalytic dwell and associated states probably accounts for the structures of F_1_ being determined primarily in this rotational position. In particular, this can be attributable to the ADP-inhibited state of F_1_, where F_1_ pauses catalysis and rotation at the catalytic dwell angle^30^, with the structure being near identical to the ground state structure^31^. To address this problem, we attempted to freeze F_1_ in the binding dwell by trapping F_1_ molecules in a temperature sensitive (TS) dwell state found in the single molecule rotation assays of TF_1_ imaged under low temperatures. These studies have shown that, when rotation of wild-type TF_1_ is observed below 10°C, the TS dwell is an intervening pause at the binding dwell angle^32^. Furthermore, the catalytic-site mutation TF_1_(β_E_190D), substantially extends the duration of the TS dwell, even at room temperatures^33^. Kinetic analysis showed that the TS dwell occurs either immediately before or after the transition from ATP-association to ATP-waiting.

Here we describe the use of cryo-Electron Microscopy (cryo-EM) to examine the temperature sensitive TF_1_(β_E_190D) mutant under three conditions, which has enabled structures representing the catalytic and binding dwell states to be obtained. The structures indicated how ATP hydrolysis opens a β subunit to induce the hydrolysis stroke in the 40° substep. This movement reorients a different β subunit that closes to bind MgATP tightly, thereby inducing the power stroke for the 80° substep. Following this step, P_i_ is released via a secondary access tunnel that allows P_i_ dissociation/association, even when bound ADP plugs the nucleotide-binding cleft. Taken together, these structures provide a molecular-level understanding on the rotary catalytic mechanism of F_1_-ATPase.

## Results

### *Structure of the Bacillus* PS3 *F_1_-ATPase in two rotational states*

Kinetic analyses from single-molecule studies estimate that at 10°C ~80% of F_1_-ATPase molecules pause at the binding dwell angle, whereas at 28°C ~85% pause at the catalytic angle^32, 33^. To obtain cryo-EM maps of F_1_-ATPase pausing at the binding dwell angle and catalytic dwell angle, MgATP was added (to a final concentration of 10 mM) to purified TF_1_(β_E_190D) that was immediately applied to EM grids at either 10°C or 28°C. These grids were then immediately frozen in liquid ethane (Extended Data Fig. 1a and b) and subsequently imaged at 300 kV followed by single particle averaging using standard sorting methods.

Cryo-EM maps were obtained to 3.1 Å and 3.4 Å resolution for the structures at 28°C and 10°C, respectively (Fig 2a and b). Although the maps were not to atomic resolution, they provided sufficient resolution to establish which nucleotide was bound in each site (Fig. 3 and Movie 1). The overall structure obtained at 28°C was almost identical to the bMF_1_ ground state; both of the β subunits were in a closed conformation and bound to MgATP, corresponding to β_TP_ or β_DP_ of bMF_1_, whereas the third β was in open conformation, representing β_E_ (Extended Data Fig. 2). Because β_DP_ represents the catalytically active state, the structure of TF_1_ obtained likely corresponds to a “hydrolysis-waiting” dwell. Although the structure was highly similar to the ground state of bMF_1_, one clear difference was found in the site occupancy of β_E_, in which MgADP was clearly present, whereas bMF_1_ does not have bound nucleotide in ground state structure.

**Figure 2:**
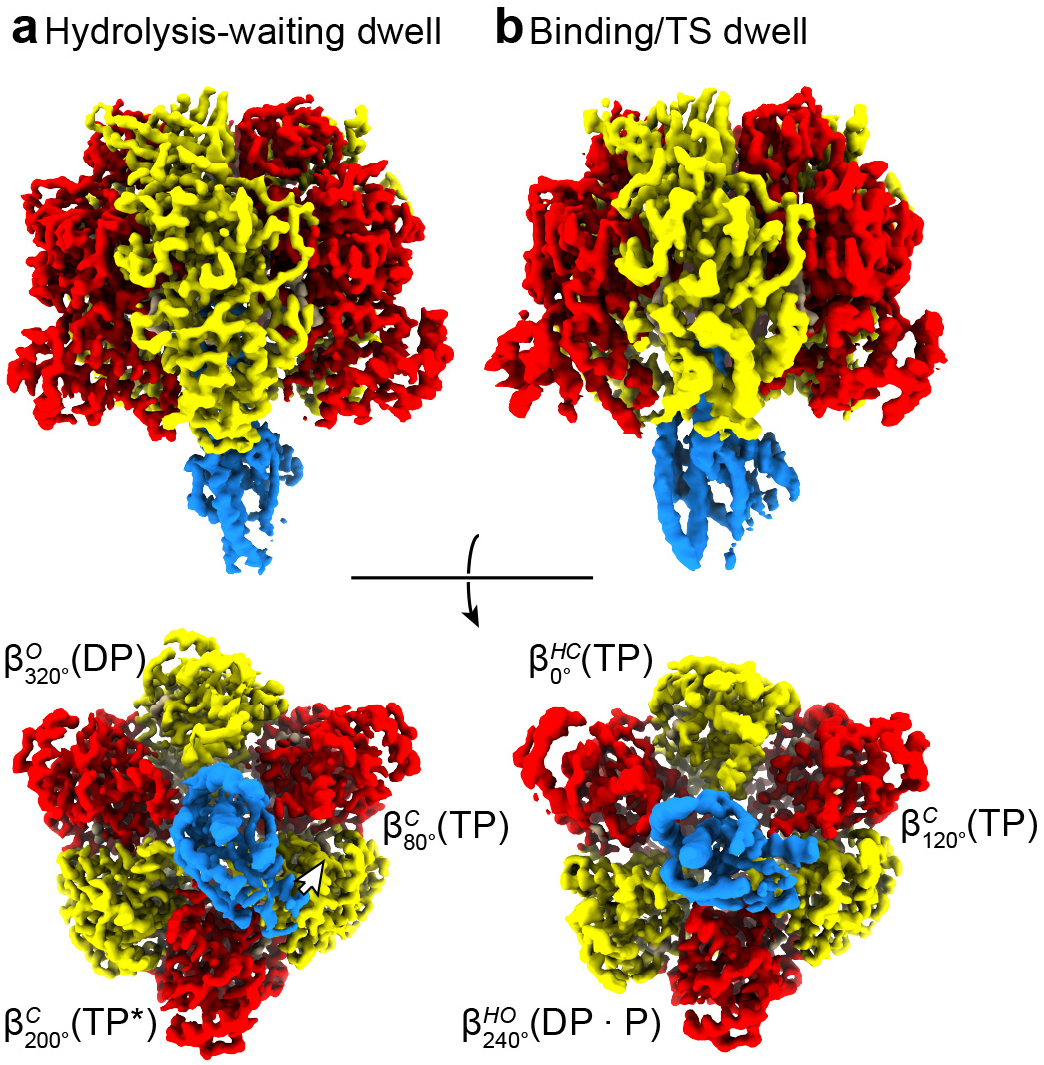
Cryo-EM maps of the *Bacillus* PS3 F_1_-ATPase βE190D in two rotational dwells. Cryo-EM maps of the two rotational dwells of TF_1_(βE190D), represented as surfaces, viewed perpendicular (top) and from the membrane (bottom), with subunit a in red, β in yellow, and γin blue. (**a**) The predominate structure when imaged following the addition of 10 mM MgATP at 28 °C – the hydrolysis-waiting dwell. (**b**) The predominate structure when imaged following the addition of 10 mM MgATP at 10 °C – the binding/TS dwell. Comparison of the binding/TS with the hydrolysis-waiting dwell indicated that between these dwells the γsubunit has rotated by ~44° in the counterclockwise direction (when viewed from the membrane – rotation highlighted with white arrow). Overall each dwell provided three different conformations of the F_1_-ATPase βsubunits and so provided a spectrum of the six sub-states through which the enzyme passes during its hydrolysis cycle, here termed 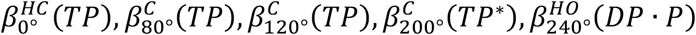 and 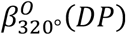.

**Figure 3:**
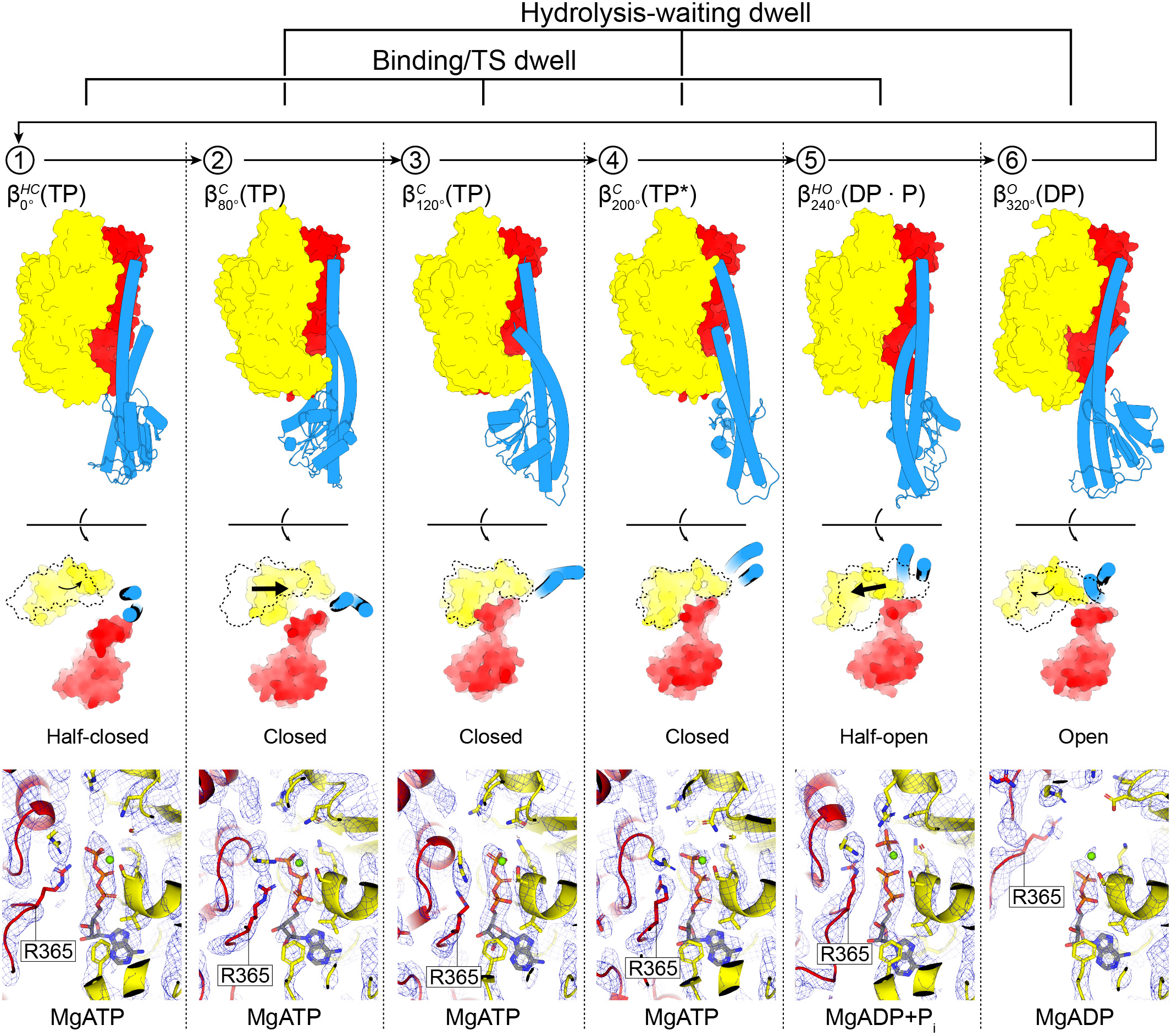
The six sequential conformations in the F_1_-ATPase βE190D rotary catalytic cycle. (**top**) αβ pairs superposed on the N-termini (β2-82) and viewed from the side (perpendicular to the membrane) and below (from the membrane). Subunit α in red, β in yellow and γ in blue, with stencil outline of the β subunit from the previous step in the scheme for comparison. (**bottom**) Close up of the catalytic nucleotide binding site, superimposed on residues around the nucleotides (β158-166, β336-342 and β412-421). Cryo-EM map shown as blue mesh. Nucleotides, Mg^2+^ and P_i_ shown as sticks with CPK coloring and αR365 (the arginine finger) labelled. In the movement from the TS dwell (states 1, 3 and 5) to the hydrolysis-waiting dwell (states 2, 4 and 6): 1 → 2 transition 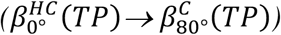, the αβ subunits close to bind MgATP tightly. 2→3→4 transition 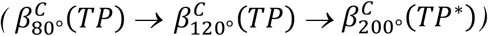 the β subunit remains in a similar position, with a minor movement of the α subunit about the nucleotide. 4→5 transition 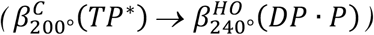, MgATP is hydrolyzed to MgADP-P_i_ and the β subunit opens to a half-open state, with the arginine finger (αR365) moving relative to the adenosine ring to stabilize the MgADP+P_i_. 5→6 transition 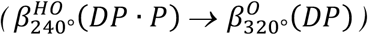, P_i_ is released and the β subunit opens. 6→1 transition 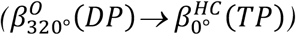 MgADP is released, followed by MgATP binding and the β subunit half closing to start the cycle again.

The structure obtained at 10°C showed features distinct from the hydrolysis-waiting dwell structure. Two of the β subunits assumed novel conformations that are intermediates between the closed and open forms, whereas the third assumed a closed conformation (Fig. 3 and 4, and Extended Data Fig. 3). Another striking feature is that the orientation of the γ subunit differed from the hydrolysis-waiting dwell structure. Fig. 2 shows the alignment on the α_3_β_3_ stator ring that gives minimum angular displacement of the γ subunit from the hydrolysis-waiting dwell structure, with the γ subunit rotated +44° from the hydrolysis-waiting dwell structure (as calculated with CCP4mg^34^). This agrees well with 40° substep from the catalytic dwell angle to the binding dwell angle observed in single molecule studies^16,33^, suggesting that the 10°C structure represents the state pausing at the binding dwell angle due to the TS dwell, as expected. Therefore, it is reasonable to consider the β subunits that correspond to β_E_, β_TP_, or β_DP_ as representing the states at 0°, 120°, and 240° in Fig. 1. All three β subunits had nucleotide bound (ATP, ATP, and ADP+P_i_, respectively - Fig. 3). This feature is also completely consistent with the expected site-occupancy of TS dwell state in Fig. 1: the state at 0° represents the state after ATP binding but before the power stroke 80° substep, the state at 120° has ATP tightly bound and the state at 240° represents the post-hydrolysis state. Therefore, we conclude that the 10°C structure represents the binding/TS dwell, pausing at the binding dwell angle, and term this the “TS dwell structure” below. The conformational states of β subunits found in the TS dwell structure are hereafter referred to as *β*_0°_, *β*_120°_, and *β*_240°_ respectively. Correspondingly, β_TP_, β_DP_, and β_E_ in the hydrolysis-waiting dwell are termed *β*_80°_, *β*_200°_, and *β*_320°_ below to ensure coherence.

**Figure 4:**
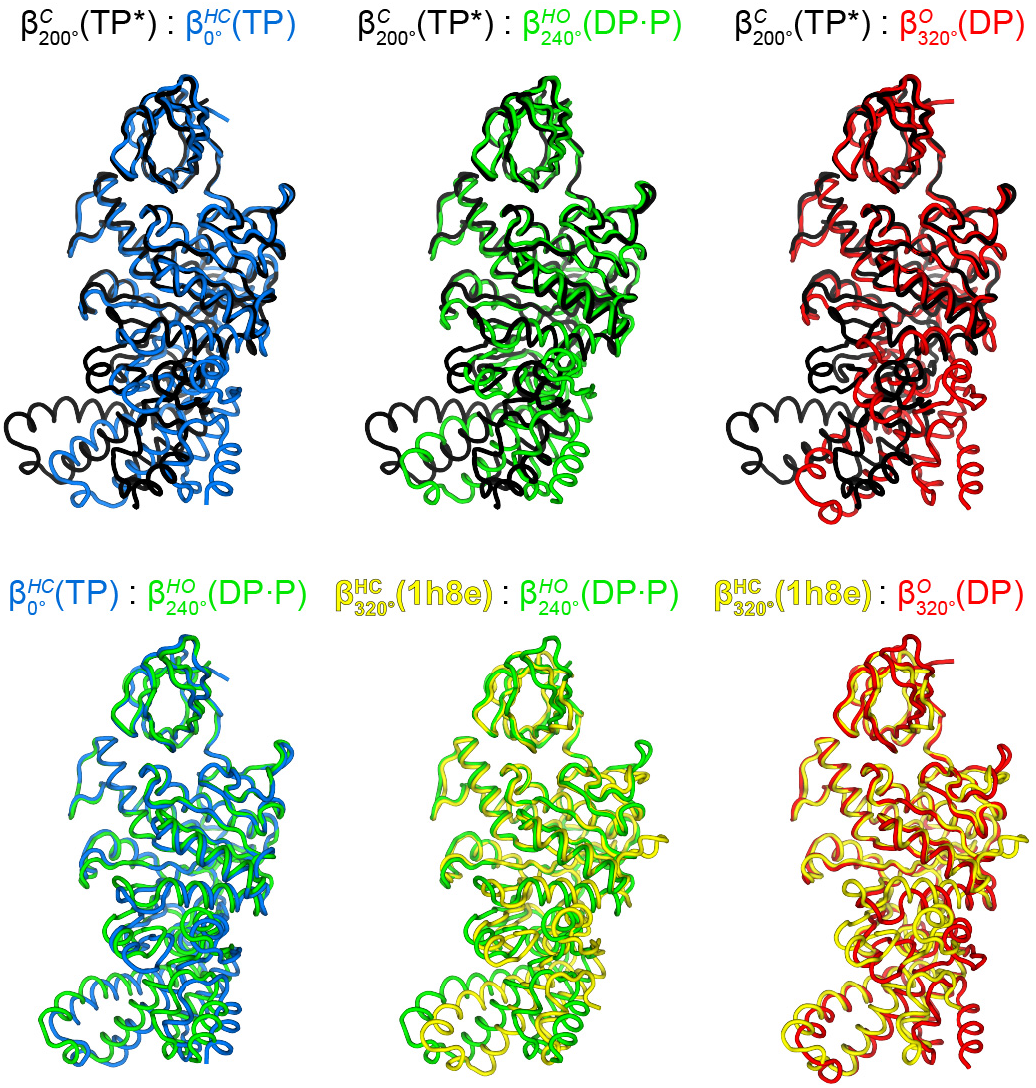
Comparison of conformational states of β subunits. Superposition (on the N-terminal β barrel) of the four main conformational states seen in the TF_1_(βE190D) (this study; 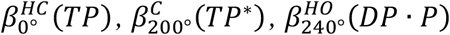 and 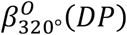) and the half open state observed for 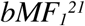 (pdb1h8e). 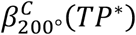 (black) is in a fully closed state, 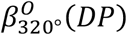 (red) is in a fully open state, 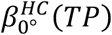 and 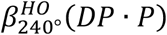 are intermediate structures that are either half closing or half opening, and 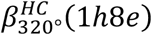 is an intermediate structure of bMF_1_ close to the open conformation. See also Extended Data Fig. 3 for a view from below.

Neither dataset contained one conformation exclusively, with the 28°C dataset containing ~3:1 and the 10°C dataset containing a ~2:3 ratio of the hydrolysis-waiting and TS dwells, respectively as classified by cryoSPARC heterogeneous refinement^35^. Although this does not agree fully with single-molecule analysis, the observed populational shift of two states well reflects the temperature-dependency of TF_1_(βE190D), suggesting that the structural analysis was valid. With the discrepancy in relative proportions possibly caused by the classification algorithm failing to sort one conformation more than the other, resulting in a greater proportion of some types of particle being discarded or particles being grouped with the minor classes. The reduced number of particles in the less abundant states resulted in poorer details in the maps, which were harder to process and consequently having the two datasets was necessary to obtain detailed information on each dwell state.

The hydrolysis-waiting dwell and TS dwell structures provide a set of the six sequential conformational states of the β subunit: *β*_0°_(*TP*), *β*_80°_(*TP*), *β*_120°_(*TP*), *β*_200°_(*TP*\*), *β*_240°_(*DP · P*), and *β*_320°_(*DP*) where the abbreviations in brackets represent bound nucleotide: *TP* for ATP, *TP*\* for ATP in catalysis, *DP · P* for ADP and phosphate, *DP* for ADP. The set of sequential conformational states are consistent with the scheme in Fig. 1, with the exception that ADP is bound to *β*_320°_. As noted above, the ground state structure of bMF_1_ did not have bound nucleotide to *β*_E_ (= *β*_320°_). An attractive interpretation is that this represents an aspect of a divergence of reaction schemes among F_1_-ATPase’s and the *β*_320°_ conformational state represents a catalytic intermediate, ADP-releasing state of TF_1_. Previous single-molecule studies showed the β subunit releases ADP during the transition from *β*_240°_ to *β*_320°_, though the exact timing was not resolved^17,18,36^ and hence ADP release may occur at *β*_320°_ or this may represent ADP re-binding from solution. However, this point remains to be addressed in future work.

Several lines of evidence indicate that P_i_ is released at the *β*_320°_ state^17,37,38^. Because F_1_ releases P_i_ within milliseconds after catalysis, it not unreasonable that we observe no bound P_i_ at the *β*_320°_ position under these conditions. However, to confirm the P_i_ binding ability of *β*_320°_, we analyzed the structure of TF_1_(βE190D) prepared in 100 mM phosphate buffer without addition of MgATP (Extended Data Fig. 1c). The structure showed essentially the same conformation as the hydrolysis-waiting dwell, except that P_i_ was bound to *β*_320°_ instead of MgADP (Extended Data Fig. 4). The P_i_-bound *β*_320°_(*P*) is consistent with the contention that P_i_ is likely released from the *β*_320°_ state. We term the resultant structure the “P_i_-bound” dwell and is in a similar rotary state to the bMF_1_ ground state, and is therefore most similar to the bMF_1_ crystal structure in the presence of Inhibitory Factor 1, AMP-PNP and thiophosphate (F_1_–ThioP)^38^. Similar to the data collected after the addition of MgATP, our analysis identified a second structure in addition to the P_i_ bound state. However this was not the binding/TS dwell, but instead a symmetrical α_3_β_3_ complex lacking the γ subunit (Extended Data Fig. 1c), which is similar to that of the TF_1_ α_3_β_3_ complex solved via crystallographic methods^39^. Close inspection of the *β*_320°_ state (in both the hydrolysis-waiting and P_i_-bound dwells) indicated the presence of an internal tunnel that passes through the central cavity of the enzyme and opens only in the *β*_320°_ state (Fig. 5). Because this tunnel has a minimum diameter of ~9 Å, it could potentially mediate the release of P_i_ when exit through the nucleotide-binding cleft was blocked by bound MgADP.

**Figure 5.**
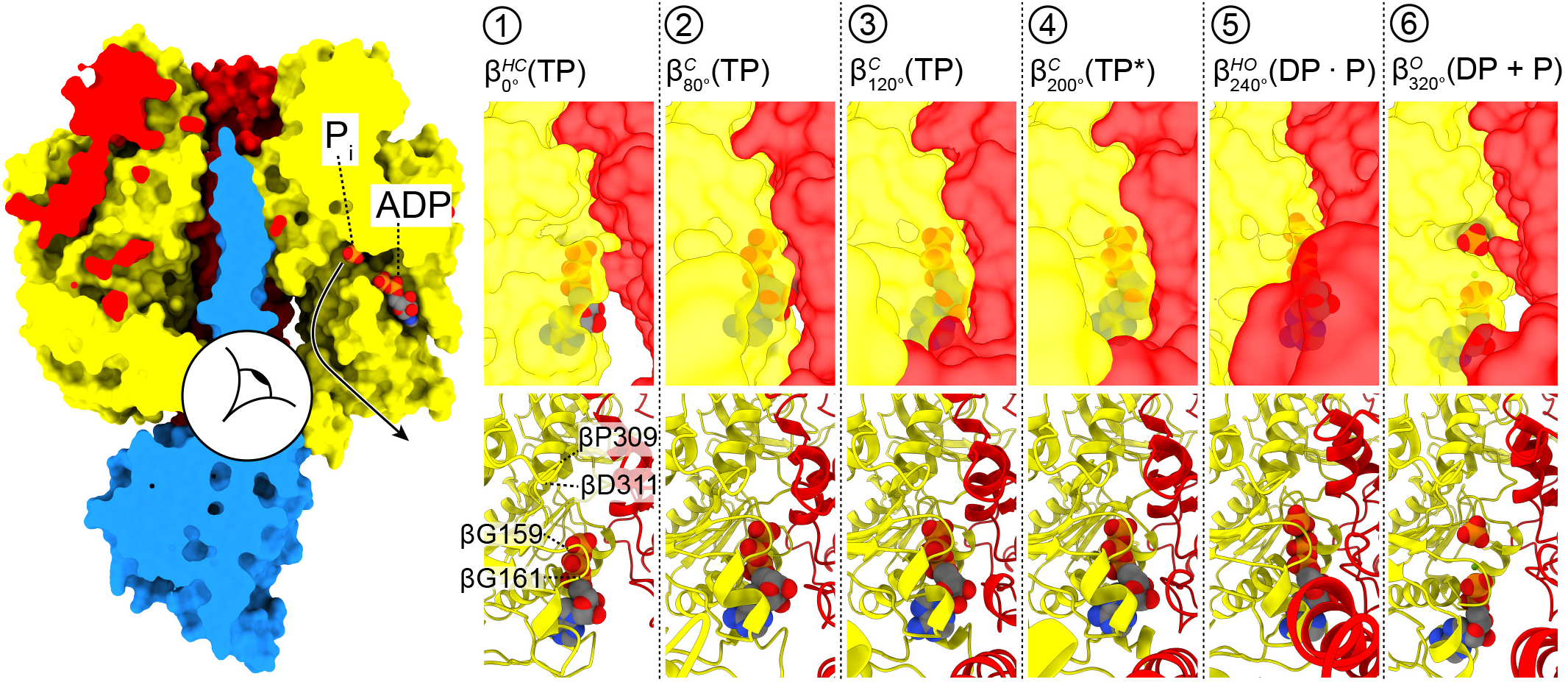
(**left**) Composite structure (to show P_i_ and MgADP bound in the same figure) of the hydrolysis-waiting and P_i_-bound dwell, cut away to observe the β_320°_ binding site with protein shown as surface and MgADP+P_i_ shown as spheres with CPK coloring. Composite made by docking the MgADP from the hydrolysis-waiting dwell into the same position in the P_i_-bound structure. The alternative P_i_ exit channel (black arrow) could facilitate dissociation of the P_i_ while MgADP remains bound. **(right)** View of the P_i_ binding site from within the complex (view depicted with eyeball in the left panel). As the enzyme cycles, the P_i_ exit channel only opens in the β_320°_ state. Loop βP309-311 closes and opens the exit channel.

The cryo-EM maps show only weak information about the sidechain at the β_E_190D substitution (Extended Data Fig. 5), which is commonly seen for carboxylate residues in cyro-EM maps^40^, and therefore it was difficult to define unequivocally how this mutation stabilizes the TS dwell. However, given its location near the Mg^2+^ ion and γ-phosphate, it likely interferes with the water network in the binding site, which could not be resolved in the present maps.

### The F_1_-ATPase rotary catalytic mechanism

The structures obtained in the present study define the complete series of conformational states assumed by a β subunit as it progresses through the F_1_-ATPase catalytic cycle (Fig. 6 and 7, Extended Data Fig. 6, and Movies 2 and 3). Starting at 0° - the TS dwell state - 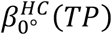 is in the half-closed conformation with MgATP bound loosely. After the TS reaction, 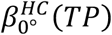 initiates the conformational transition from half-closed to closed form 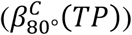 through a typical induced-fit conformational transition, driven by the progressive development of the affinity between the enzyme and bound MgATP, the so called ‘binding-change’. This accompanies the pull-motion of the C-terminal domain of the β subunit that induces the 80° substep rotation of the γ subunit. The contribution of the binding-change process for torque generation is estimated to be 21-54 pNnm^41^, and so is the major force-generating step. Throughout the rotation from 80° to 200°, the β subunit remains in the closed form. At 200°, while the β itself is still in closed form, the α/β interface becomes closed. As a result, the catalytically critical arginine residue of the α subunit (αArg364 in TF_1_), commonly called the *arginine-finger*, has contact with the γ phosphate of ATP that triggers ATP hydrolysis. Following this substep, the β subunit executes a second conformational transition - from the closed form to a half-closed form - resulting in generation of the 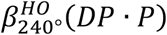 substate, as also suggested in a previous single-molecule study^42^. At 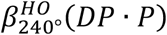, the posthydrolysis state (ADP+Pi) is stabilized. This conformational change results in a shift in the equilibrium to favor hydrolysis and thereby contributes to torque generation. However, this contribution is estimated to be only 7-17pNnm^41^, somewhat lower than that from the binding-change process. Subsequently, upon rotating from 240° to 320°, the β subunit transforms to fully open conformation to generate 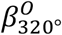. Although this third conformational transition likely generates torque as well, the extent of its contribution has not yet been investigated, but is likely small due to the majority of torque being accounted for by the power stroke. We observed two states at the 320° position 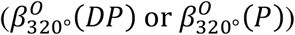. Although the present results do not allow us to identify unequivocally which state represents the catalytic intermediate, singlemolecule studies support the latter^17,36,37^. Fig. 5 shows both ADP and P_i_ present at the 320° rotary position and intriguingly we observed an alternative exit pathway (tunnel) for P_i_ release. After releasing ADP/P_i_, the β subunit then returns to the starting position at 0°. The conformational state of *β*_0°_ before it binds MgATP is not clear, however, because the nucleotide-free β preferentially takes an open conformation in the isolated α_3_β_3_ subcomplex, it is reasonable to assume that *β*_0°_ is in an open conformation before ATP binds. Then, upon ATP association, 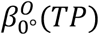 transforms to a half-closed conformation, 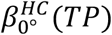. This conformational transition may not accompany the rotation of the γ subunit.

**Fig. 6.**
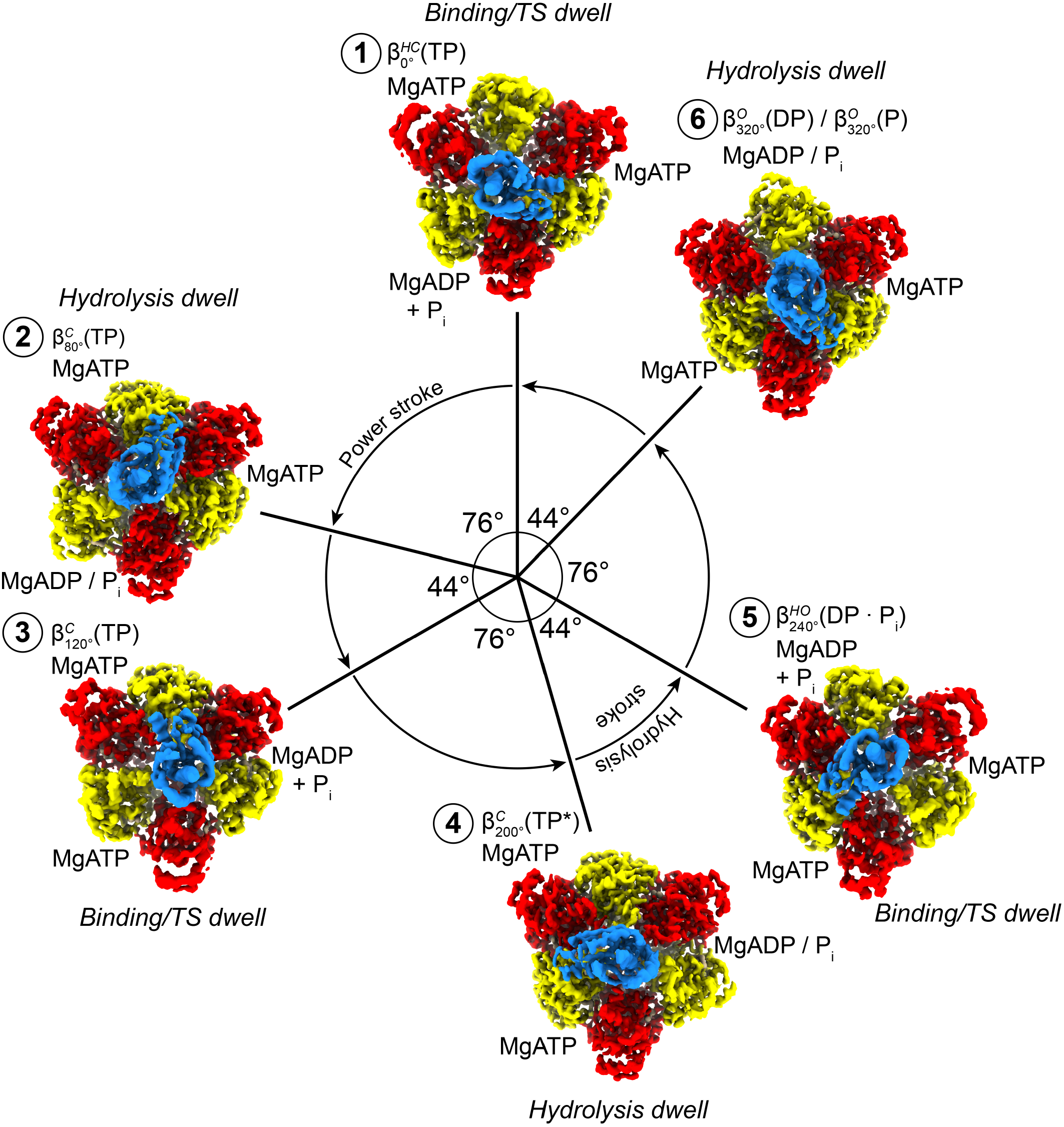
The reaction scheme shown as a wheel diagram. Circle with arrows describes rotation of the γ subunit. Cryo-EM maps for each state shown as surfaces, with subunit α in red, β in yellow and γ in blue. ATP is sequentially bound and hydrolyzed, causing the power stroke (76°rotation) and hydrolysis stroke (44 °rotation).

**Figure 7:**
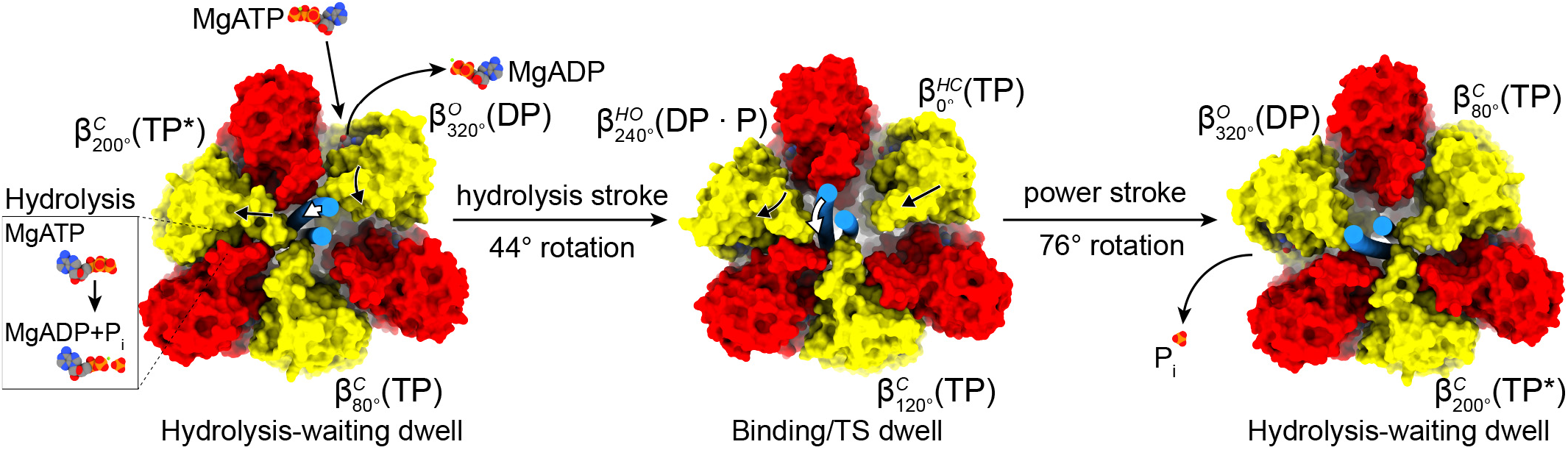
Torque generation in the F_1_-ATPase. The 120 ° rotation of the γ subunit is achieved by two successive rotations. Subunits α and β are shown as surface, the coiled coil section of subunit γ as two cylinders and nucleotide/P_i_ as spheres with CPK coloring. The hydrolysis stroke occurs after the hydrolysis-waiting dwell; MgATP is hydrolyzed to MgADP+P_i_ in the 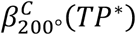 site resulting in its opening to a half-open state, so that the γ subunit is pulled towards the 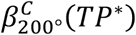 site and pivots around the β_TP1_ subunit with a 44° rotation, MgADP is exchanged for MgATP in the 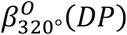 site which reorients to a half-closed state primed for the power stroke. The power stroke occurs after the TS dwell; MgATP is tightly bound by the 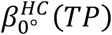 site, closing to a closed state, and 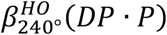 opens to and open state, with the γ subunit being is pushed towards the 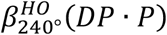 site, pivoting 76° around the 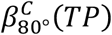 subunit. P_i_ is released at the hydrolysis-waiting dwell once the P_i_ exit tunnel is open.

The two principal movements are the power stroke (after the TS dwell), which is driven by the tighter binding to MgATP, and the hydrolysis step (following the hydrolysis-waiting dwell), with the associated conformational changes being transferred to the γ subunit to induce its rotation (Fig. 7). An interesting feature observed in this study is that the closing of the open subunit to the half-closed state reorients the β subunit around the γ subunit leaving it in a mechanically favorable position to perform the power stroke (Fig. 7 and Movie 2). This is enabled through the different trajectories that the β subunit closes as it transitions from the open to the half closed and then closed state (Movie 3). The difference in trajectory can be observed by superposition of the β subunits on the N-terminal β-barrel and defining the rotation axis of the “foot” of the β subunit (residues 129-180 and 327-470) (Extended Data Fig. 7). The difference in the rotation axes results in the foot either pushing towards or away from the γ subunit (open ^®^ half closed) or twisting around the γ subunit (half closed ^®^ closed), with angle between these axes being ~60°. A further key observation is that the rotation of the γ subunit during catalysis pivots about the other β, which remains stationary and tightly bound to MgATP (Fig. 7). Notably, the axis about which these two rotations occurs wobbles (or precesses) during rotation, resulting in a change in the radius of the γ subunit rotation between the power stroke and hydrolysis events (Extended Data Figure 8) that has also been observed in single-molecule studies^43^.

## Discussion

Using cryo-EM it has been possible to observe the *Bacillus* PS3 F_1_-ATPase rotating through six sub-states and correlate the structural changes as the enzyme progresses through its catalytic cycle with single molecule experiments. Cryo-EM has the advantage of avoiding the influence of crystal lattice contacts on the position of the γ subunit observed in X-ray crystallography^1,44–46^, and has enabled detailed information to be obtained on the ways in which MgATP binding and hydrolysis generate conformational changes in the F_1_-ATPase that result in the rotation of the γ subunit. These studies have also highlighted a previously unidentified tunnel that could mediate P_i_ release while the nucleotide-binding cleft remained blocked by MgADP.

The structures obtained here provide a complete rotary catalysis model for TF_1_(βE190D), showing how MgATP is first tightly bound, resulting in the power stroke, and then cleaved, resulting in the hydrolysis stroke (Fig. 7 and Extended Data Fig. 9). The residues around the adenosine ring do not change substantially between any of the states, consistent with single molecule studies on the F_1_-ATPase using a base-free triphosphate^47^. Moreover, our results also indicate that P_i_ can be released, even when the nucleotide-binding cleft is obstructed by ADP, through an alternative exit channel, which likely explains why the affinity for P_i_ was strongly dependent on the angle of rotation^37,48^, with ATP binding likely being prevented while the lysine of the Walker A motif (βK164) is coordinated to the P_i_ (Extended Data Fig. 4). An alternate pathway for P_i_ release has been suggested previously using molecular dynamic simmulations^49^, but the energy required to enable exit was still large because the tunnel was closed in the structures used for these calculations. The alternate exit path we define here is akin to the ‘back door’ model that has been suggested to be important for force generation in myosin^50^, highlighting a potential conserved mechanism between the motors. During ATP synthesis, the ability to bind nucleotide and P_i_ through different pathways would be beneficial preventing the system being locked if MgADP binds first.

### Data availability

The models generated and analyzed during the current study are available from the RCSB PDB: 7L1Q, 7L1R and 7L1S. The cryo-EM maps used to generate models are available from the EMDB: 23115, 23116 and 7L1S.

## Acknowledgments

We wish to thank and acknowledge Dr Simon Brown for aiding in data collection and the use of the University of Wollongong Cryogenic Electron Microscopy Facility at Molecular Horizons under the management of Dr. James C. Bouwer and Directorship of Dr. Antoine van Oijen. We also wish to thank and acknowledge Dr Sawako Enoki for providing data of kinetics of TF_1_(βE190D) and Mr. Yi Zeng for cryo-EM data processing advice. We also wish to thank and acknowledge the use of the Victor Chang Innovation Centre, funded by the NSW Government, and the Electron Microscope Unit at UNSW Sydney, funded in part by the NSW Government. Molecular graphics and analyses performed with UCSF ChimeraX, developed by the Resource for Biocomputing, Visualization, and Informatics at the University of California, San Francisco, with support from National Institutes of Health R01-GM129325 and the Office of Cyber Infrastructure and Computational Biology, National Institute of Allergy and Infectious Diseases. A.G.S was supported by a National Health and Medical Research Council Fellowship APP1159347 and Grant APP1146403. This work was supported in part by Grant-in-Aid for Scientific Research on Innovation Areas (JP18H04817, JP19H05380) from the Japan Society for the Promotion of Science.

## Contributions

M.S. performed the formal analysis of the study. H.U. revised the manuscript and provided the clones and purification strategy. H.N. and A.G.S. conceived and supervised the study and drafted the manuscript.

## Competing interests

Authors declare no competing interests

## Methods

### Protein purification

JM103Δunc *E. coli* cells harboring the expression plasmid for PS3 F_1_-ATPase (HCXP) βE190D were incubated at 37°C with 170 rpm shaking overnight (~18 h) in a baffled flask containing 1 L Terrific broth with 100 μg/mL carbenicillin. Cells were collected at 4,000 g resulting in ~7.5 g of cells. The cell pellet was resuspended in 75 ml of 50 mM imidazole, 100 mM NaCl (pH7.0), with one cOmplete EDTA-free Protease Inhibitor Cocktail Tablet (Roche) and DNase1 at 4°C. Cells were lysed with sonication for 3 min at 4°C, and cell debris removed by centrifugation at 50,000g for 40 min at 4°C. The supernatant was then applied to a 3 ml gravity flow Ni-NTA column that had been pre-equilibrated in 50 mM imidazole and 100 mM NaCl (pH7.0). The column was washed with 40 column volumes of 75 mM imidazole 100 mM and NaCl (pH7.0) and eluted with 5 column volumes of 500 mM imidazole and 100 mM NaCl (pH7.0). Fractions containing F_1_-ATPase (as assessed by SDS PAGE) were pooled and concentrated to 550 μL, before application to a Superdex 200 10/300 GL column (GE Healthcare) equilibrated in 100 mM Potassium phosphate buffer and 2 mM EDTA (pH7.0). Fractions containing F_1_-ATPase (as assessed by SDS PAGE) were pooled and concentrated to 8 mg/ml (300 μL).

### Cryo-EM grid preparation

For the Phosphate buffer experiment, 3.5 μl of purified protein was transferred to a glow-discharged holey gold grid (Ultrafoils R1.2/1.3, 200 Mesh). Grids were blotted for 4 s at 22°C, 100% humidity, and flash-frozen in liquid ethane using a FEI Vitrobot Mark IV. For the +10 mM MgATP at 28°C or 10°C experiments, 30 μL of protein was buffer exchanged into 20 mM Tris (pH7.0) 50 mM KCl using a Amicon Pro spin column. 4.5 μl of protein was then incubated at 28°C or 10°C in a thermocycler for 20 min. 0.5 μL of 100 mM MgATP (at either 28°C or 10°C) was added to the protein sample and vigorously mixed with a pipette before 3.5 μl was transferred to a glow-discharged holey gold grid (Ultrafoils R1.2/1.3, 200 Mesh). Grids were blotted for 4s at 28°C or 10°C, 100% humidity and flash-frozen in liquid ethane using a FEI Vitrobot Mark IV (total time from addition of MgATP to freezing was ~15 s).

### Data collection

Grids were transferred to a Thermo Fisher Talos Arctica transmission electron microscope (TEM) operating at 200 kV and screened for ice thickness and particle density. Grids were subsequently transferred to a Thermo Fisher Titan Krios TEM operating at 300 kV equipped with a Gatan BioQuantum energy filter (with 40 eV slit) and K2 Camera. Images were recorded with tilt angles of 20-30° automatically using EPU with 60,000X magnification (microscope user interface listed magnification of 165,000X due to the energy filter) yielding a pixel size of 0.84 Å. A total dose of 50 electrons per Å^2^ was used and spread over 40 frames, with a total exposure time of 5.0 s. 3,156, 2,965 and 2,158 movie micrographs were collected for the 28°C, 10°C and phosphate buffer datasets, respectively (Extended Data Fig.1).

### Data processing

cryoSPARC^35^ was used to perform all image processing and refinement. Micrographs were first motion corrected and defocus was estimated using patches. Particles were automatically picked using cryoSPARC. *Ab initio* and heterogenous refinement were used to 3D classify the particles into multiple states (see Extended Data Fig. 1). These states were then fully refined using homogenous refinement. DeepEMhancer^51^ was used to create maps that were easy to interpret for the figures showing the entire complex (e.g. continuous density for the γ subunit, which was lower resolution than the rest of the map). Extended Data Table 1 shows data collection and refinement statistics, Extended Data Fig. 10 contains FSC curves and Extended Data Fig. 11 provides local resolution estimates.

### Model building

Models were built and refined in Coot^52^, PHENIX^53^ and ISOLDE^54^ using pdbs 6N2Y^27^ (*Bacillus* PS3 F_1_F_o_ cryo-EM structure) and 4XD7^26^ (*Bacillus* PS3 F_1_-ATPase crystal structure) as guides. Extended Data Table 1 for refinement and validation statistics.

## SUPPLEMENTARY INFORMATION

**Extended Data Figure 1:**
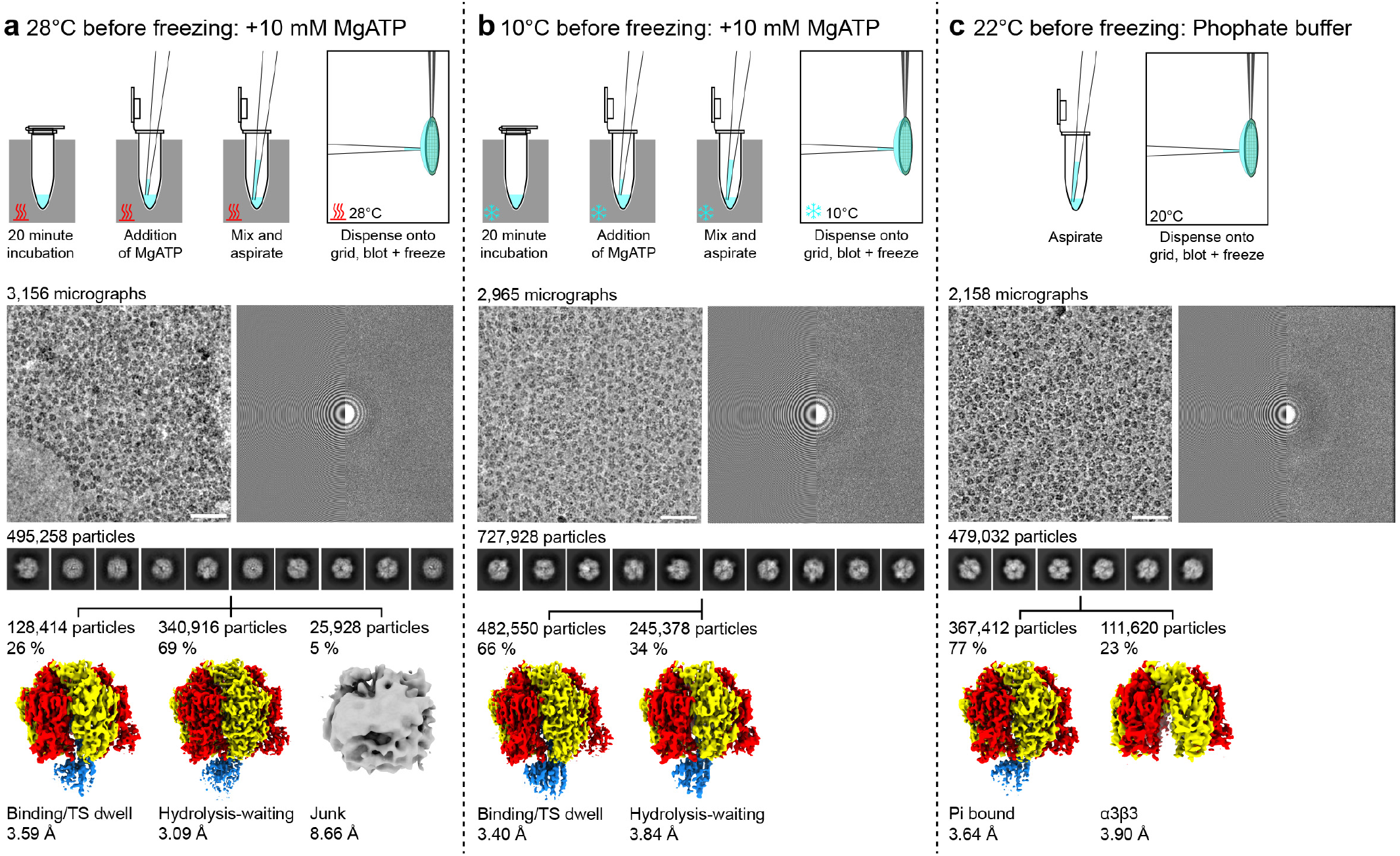
Sample preparation, data collection and data processing. (**a**) 28°C and 10 mM MgATP prior to plunge freezing. (**b**) 10°C and 10 mM MgATP prior to plunge freezing. (**c**) 22°C in phosphate buffer prior to freezing. Top: schematic of sample preparation. A thermocycler was used to heat or cool the sample prior to freezing and a Vitrobot (Thermofisher) was used to control the temperature of the grid prior to plunging freezing. Middle: sample electron micrograph (white scale bar is equivalent to 50 nm) and power spectrum of sample. Bottom: processing flowchart showing 2D classification and heterogeneous sorting (run in cryoSPARC^35^).

**Extended Data Figure 2:**
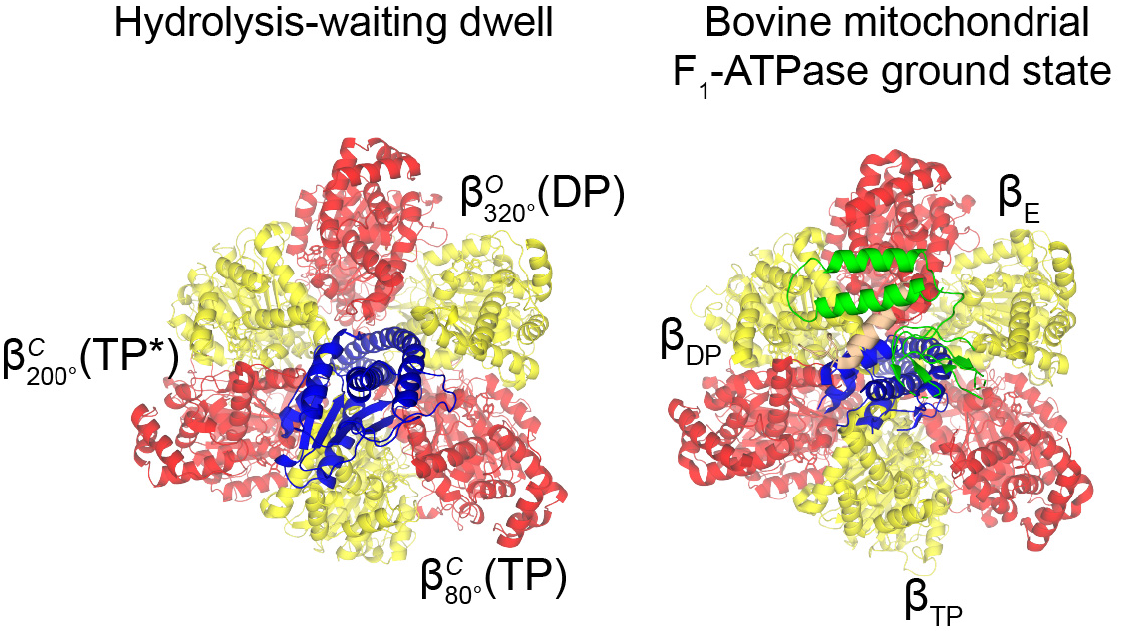
Comparison of the hydrolysis-waiting dwell with the bovine mitochondrial F_1_-ATPase ground state. Superposed structures of the hydrolysis-waiting dwell from this study with the bovine mitochondrial bMF_1_ ground state (pdb2JDI^19^) shows that they are in a similar rotational dwell/state.

**Extended Data Figure 3:**
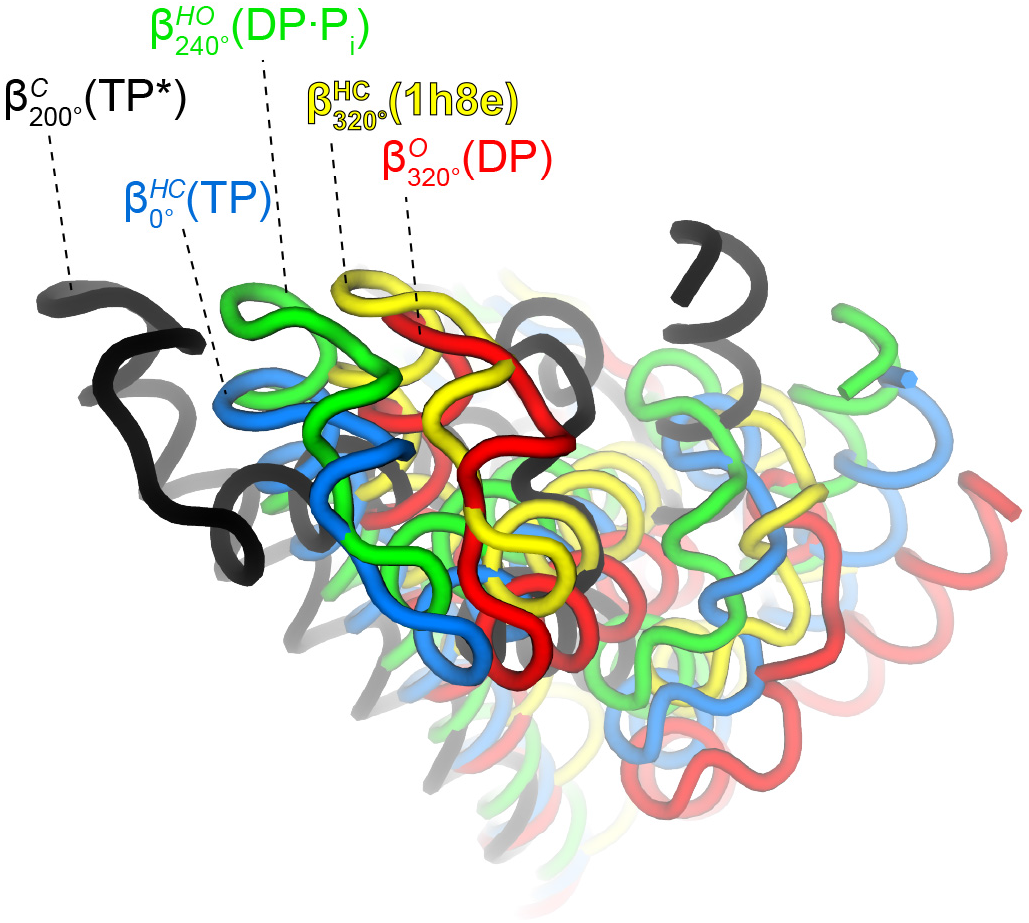
Comparison of conformational states of β subunits, viewed from below. Figure to show the relative conformations of the β subunits. Same colors and superposition as Figure 4, but combined and viewed from below (from the membrane).

**Extended Data Figure 4:**
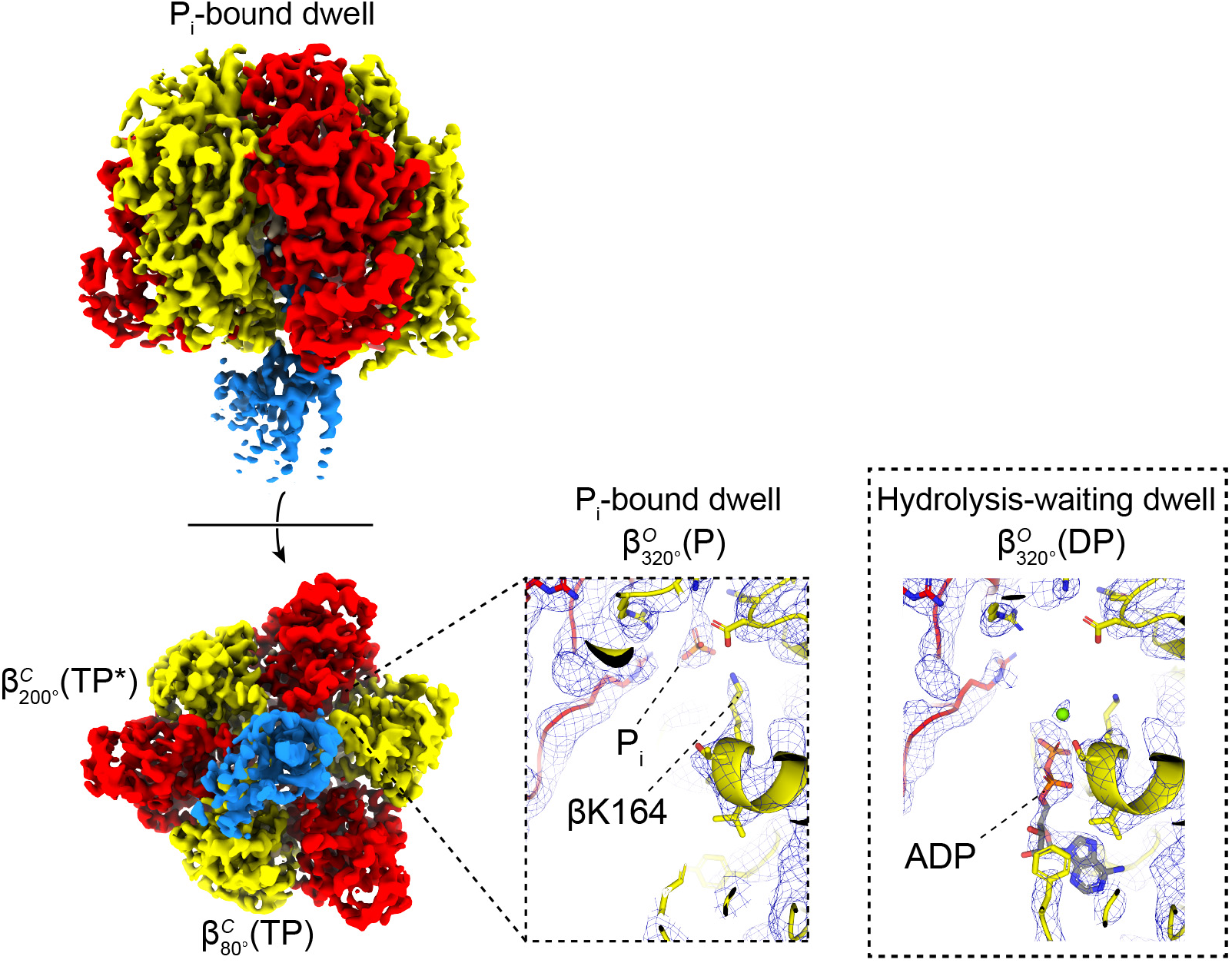
Cryo-EM structure of the P_i_-bound dwell. The P_i_-bound dwell is in the same rotational state as the hydrolysis-waiting dwell and contains P_i_ in the 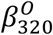 site. Cryo-EM map shown as surface viewed perpendicular and from the membrane. Subunit α in red, β in yellow and γ blue. Views of the nucleotide binding sites of the P_i_-bound and hydrolysis-waiting dwells shown for comparison; cryo-EM map shown as blue mesh, and the atomic model shown as cartoon and sticks with CPK coloring (same view as Fig. 2).

**Extended Data Figure 5:**
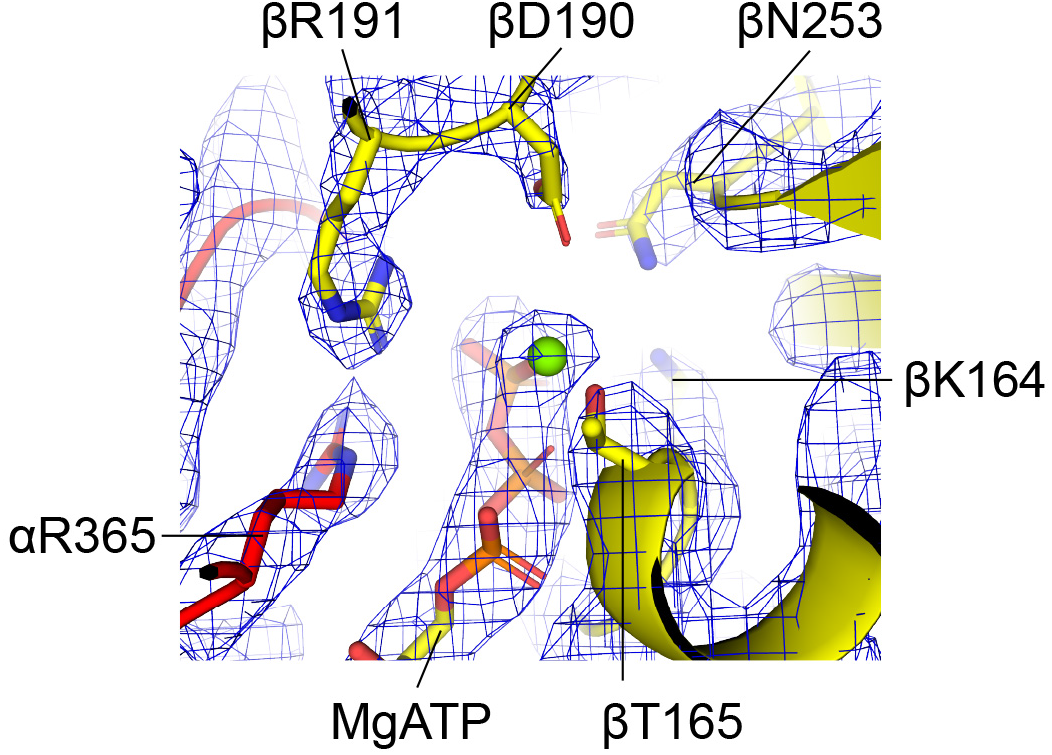
The βE190D mutation. Close up of the βE190D mutation in the 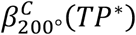 state shows that the aspartate is positioned close to the Mg^2+^ ion.

**Extended Data Figure 6:**
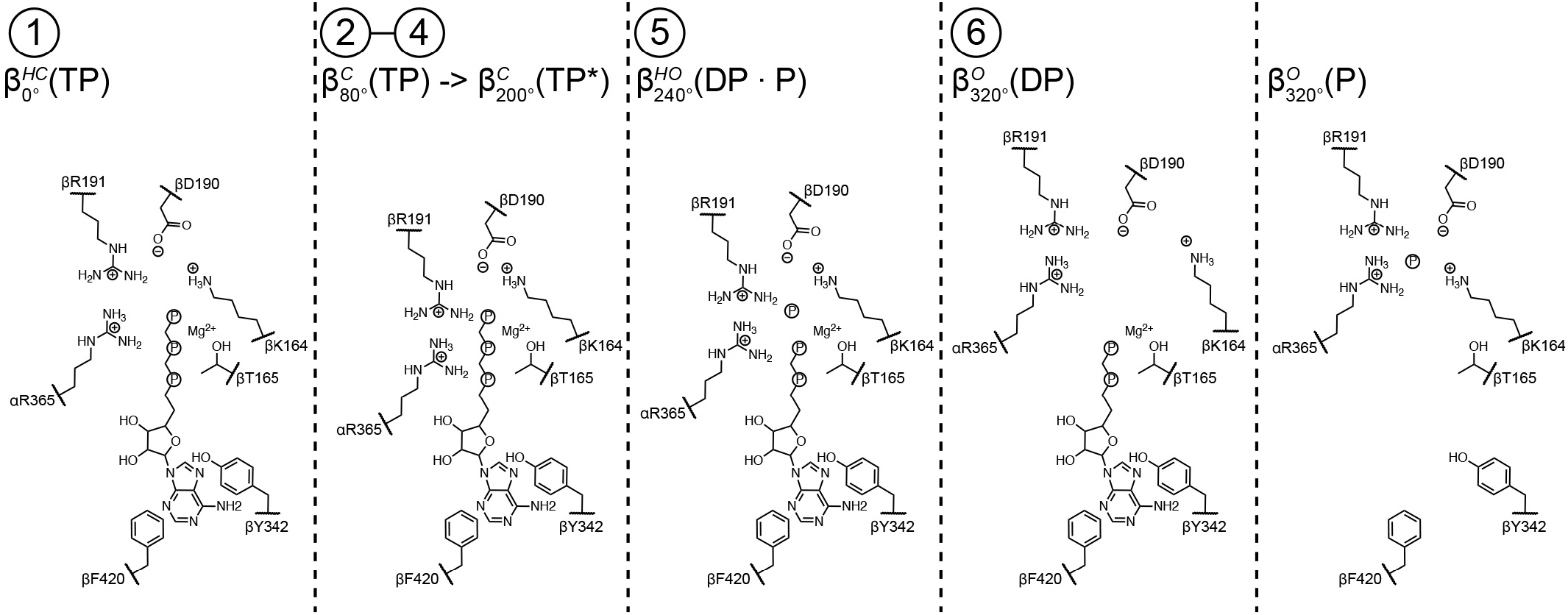
Schematics of the β nucleotide binding sites. Simplified chemical representations of the catalytic binding sites, made with ChemDraw (PerkinElmer). βY342 and βF420 hold the adenosine ring of ATP. ATP binding causes αR365, βK164 and βR191 to close onto the g-phosphate. ATP hydrolysis liberates the g-phosphate which is coordinated by αR365, βK164 and βR191. Phosphates are shown as the letter P within a circle without oxygens to aid clarity.

**Extended Data Figure 7:**
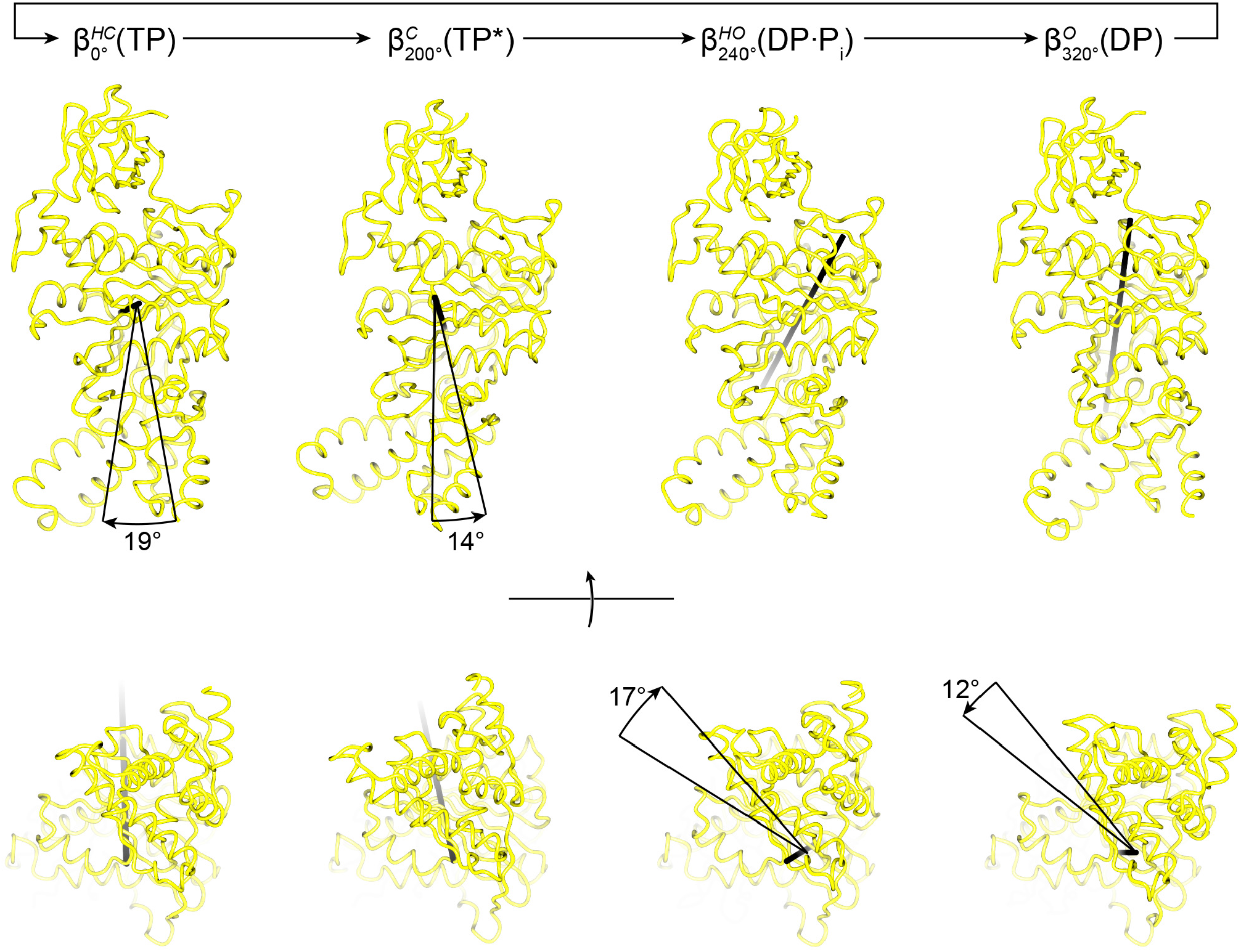
The transitions from open to closed and closed to open have two movements with different rotation axes. β subunits superposed on the N-terminal β barrel, and rotation axes of the rigid bbody defined by residues 129-180 and 327-470 shown as a black vector. Rotation axes and rotations angels calculated with ccp4mg^34^. The angle between the rotation axes for open→half closed and half closed→closed is 59°. The angle between rotation axes for closed→half open and half open→open is 50°.

**Extended Data Figure 8:**
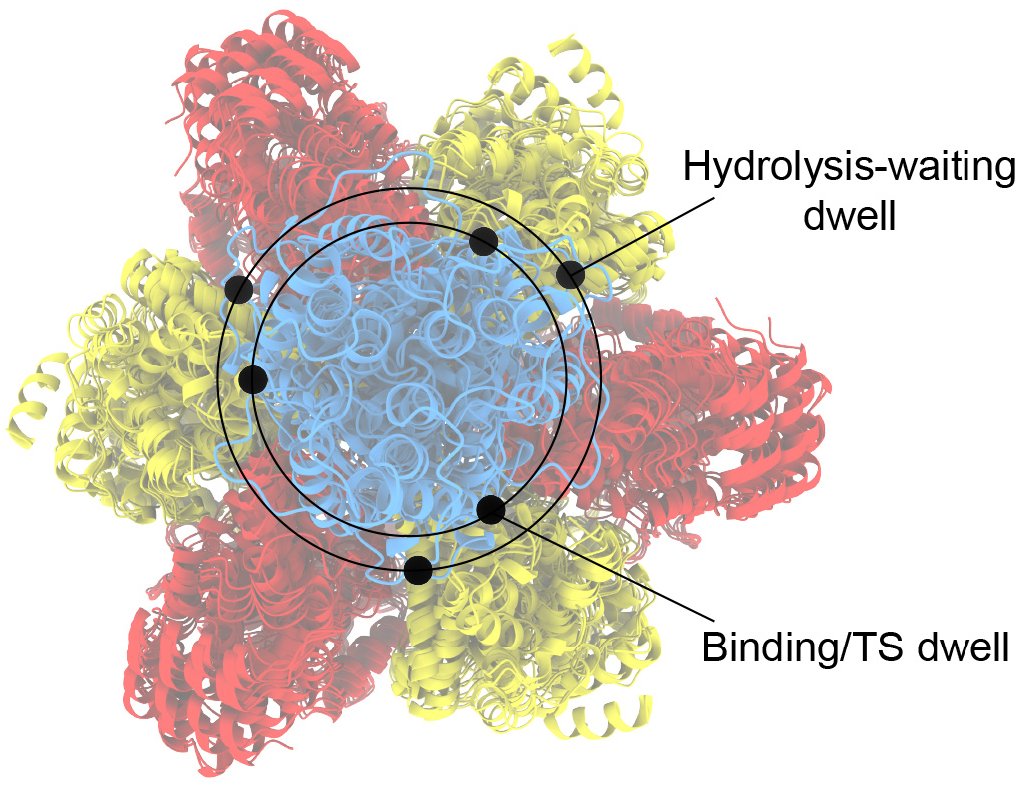
The γ subunit precesses or wobbles as it rotates. Superposition of the six rotational positions (on residues β2-82) of the enzyme shows that the two dwell states have different rotation radii. Protein shown as cartoon with black spheres for the Cα of γC112.

**Extended Data Figure 9:**
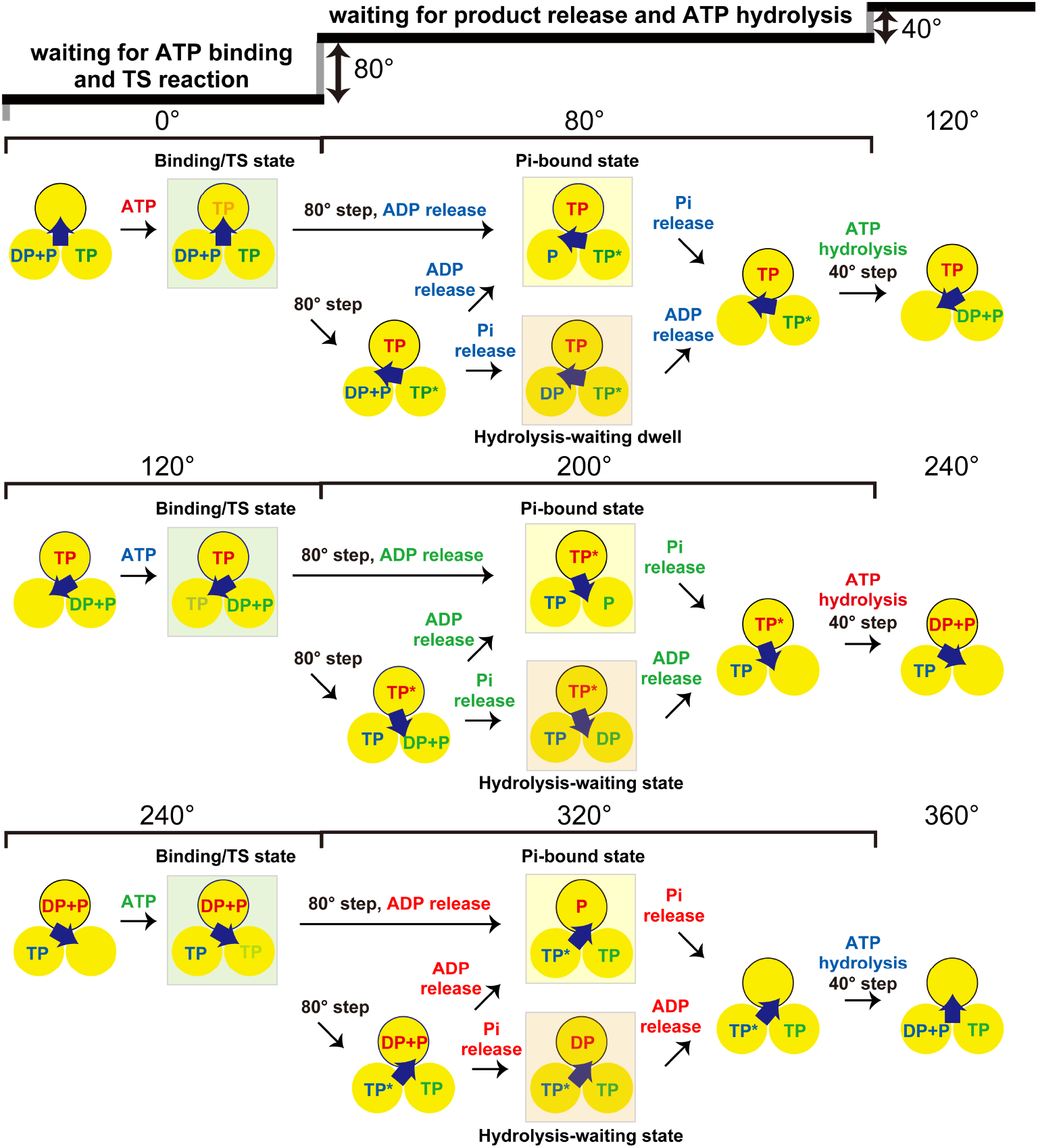
Possible models of chemo-mechanical coupling of TF_1_ (βE190D). Yellow circles represent the chemical states of catalytic sites in β subunits. The central blue arrows represent the orientations of γ subunits. TP, DP and P indicate ATP, ADP and Pi, respectively. TP* indicates ATP in catalysis. Green, yellow and orange squares indicate the states considered from the cryo-EM structures obtained in this study. 0° is defined as the ATP binding angle for the catalytic site outlined in black. In these models, ATP (red) bound at 0° is hydrolyzed to ADP and P_i_ at 200°. Three possible schemes are shown for product release. (i) ADP (red) is released during the 80° substep from 240° to 320° and then P_i_ is released at 320°. (ii) ADP (red) is released at 320° after the 80° substep from 240° to 320° and then P_i_ is released at 320°. (iii) P_i_ (red) is released at 320° after the 80° substep from 240° to 320° and then ADP (red) is released at 320°. Other catalytic sites also obey the same reaction scheme offset by 120° and 240°.

**Extended Data Figure 10:**
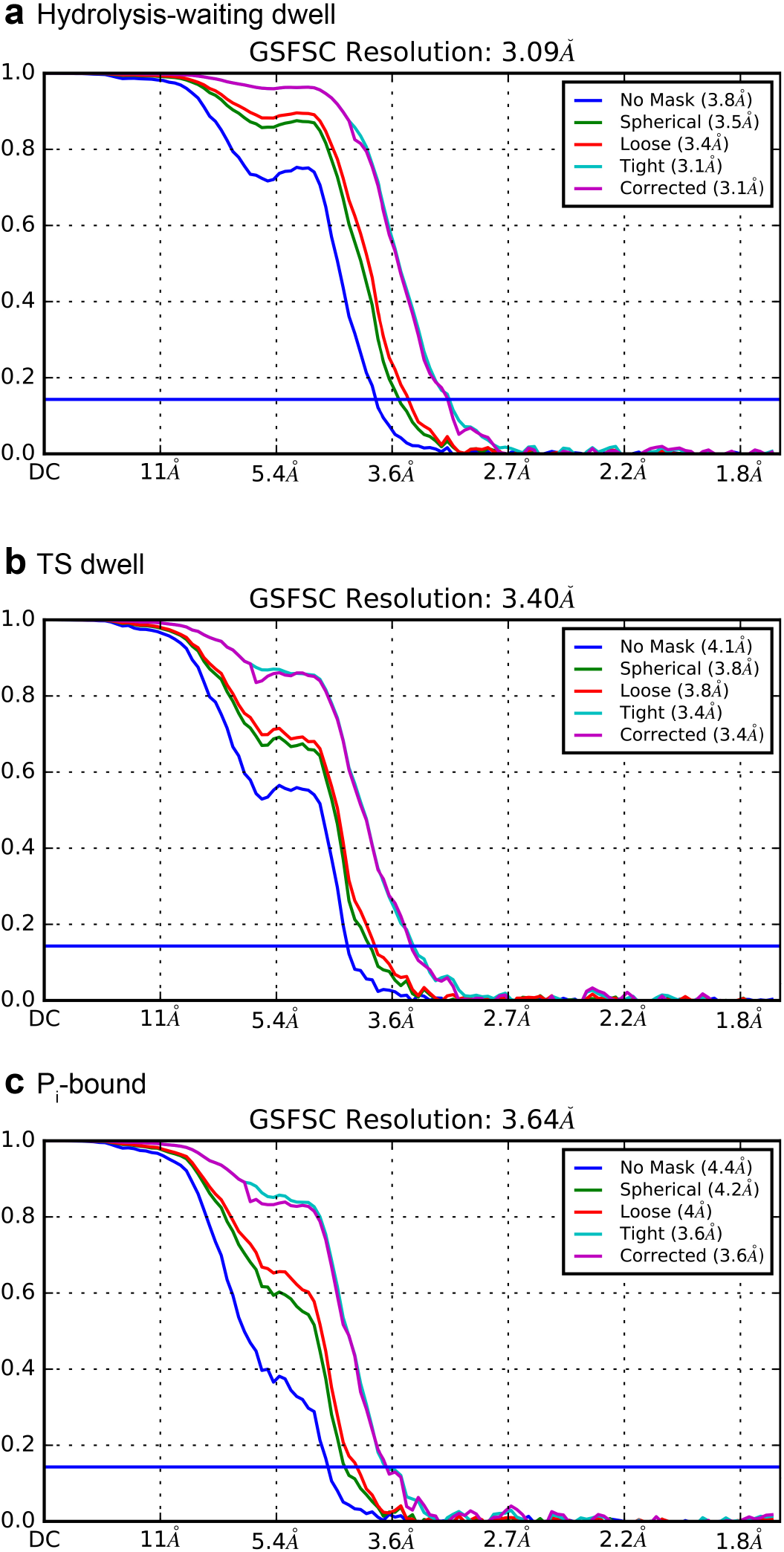
FSC curves. FSC outputs from cryoSPARC^35^. (**a**) hydrolysis-waiting, (**b**) TS dwell and (c) Pi-bound state.

**Extended Data Figure 11:**
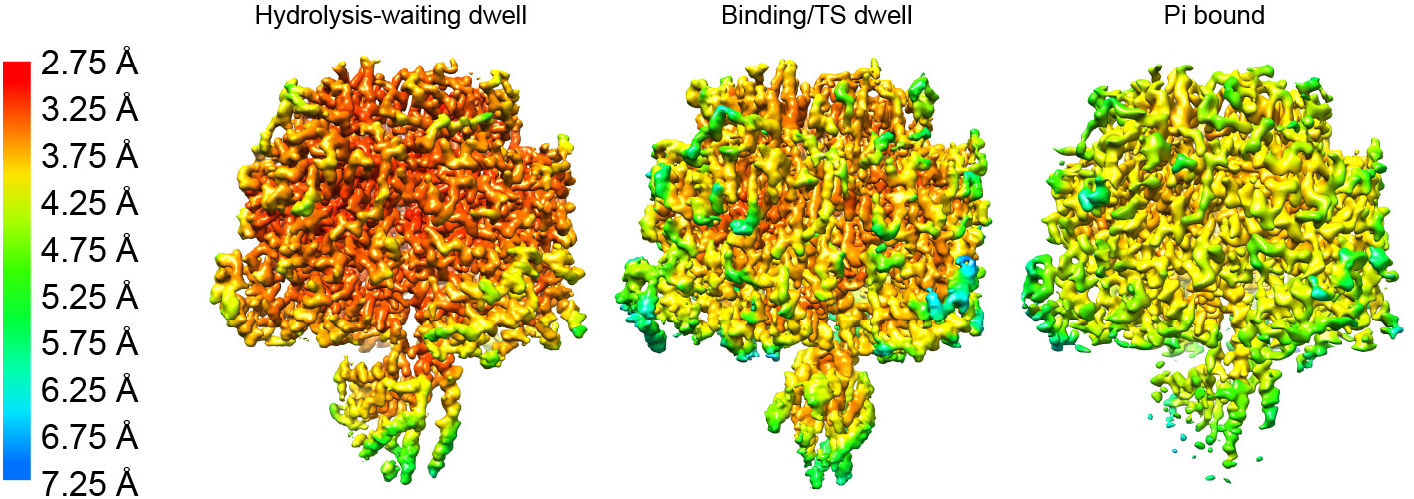
Local resolution estimates. Local resolution plots for the hydrolysis-waiting, binding/TS and Pi bound cyro-EM maps. Implemented in cryoSPARC^35^ and displayed using UCSF Chimera^55^.

**Extended Data Table 1:** Cryo-EM data collection, refinement and validation statistics.

|  | #1 Hydrolysis-waiting<br>dwell<br>(EMDB-23116)<br>(PDB 7L1R) | #2 Binding/TS dwell<br>(EMDB-23115)<br>(PDB 7L1Q) | #3 P<sub>i</sub>-bound dwell<br>(EMDB-23117)<br>(PDB 7L1S) |
| --- | --- | --- | --- |
| **Data collection and processing** |  |  |  |
| Magnification | 60,000 | 60,000 | 60,000 |
| Voltage (kV) | 300 | 300 | 300 |
| Electron exposure (e<sup>-</sup>/Å<sup>2</sup>) | 50 | 50 | 50 |
| Defocus range (μm) | 0.5-1.5 | 0.5-1.5 | 0.5-1.5 |
| Pixel size (Å) | 0.84 | 0.84 | 0.84 |
| Symmetry imposed | C1 | C1 | C1 |
| Initial particle images (no.) | 495,258 | 727,928 | 479,032 |
| Final particle images (no.) | 340,916 | 482,550 | 367,412 |
| Map resolution (Å) | 3.1 | 3.4 | 3.6 |
| 0.143 FSC threshold |  |  |  |
| **Refinement** |  |  |  |
| Initial models used (PDB codes) | 6N2Y, 4XD7 | #1 Hydrolysis-waiting dwell | #1 Hydrolysis-waiting dwell |
| Model resolution (Å) | 3.2/3.2 | 3.9/3.9 | 4.0/4.3 |
| 0.5 FSC threshold, masked/unmasked |  |  |  |
| Map sharpening *B* factor (Å<sup>2</sup>) | 108 | 115 | 146 |
| Model composition |  |  |  |
| Non-hydrogen atoms | 24,199 | 24,204 | 24,176 |
| Protein residues | 3,118 | 3,118 | 3,118 |
| Ligands | 12 | 13 | 11 |
| *B* factors (Å<sup>2</sup>) |  |  |  |
| Protein | 161 | 68 | 161 |
| Ligand | 152 | 69 | 145 |
| R.m.s. deviations |  |  |  |
| Bond lengths (Å) | 0.012 | 0.008 | 0.012 |
| Bond angles (°) | 1.784 | 1.278 | 1.832 |
| Validation |  |  |  |
| MolProbity score | 1.78 | 1.63 | 1.90 |
| Clashscore | 3.87 | 2.65 | 4.37 |
| Poor rotamers (%) | 3.32 | 2.58 | 4.07 |
| Ramachandran plot |  |  |  |
| Favored (%) | 96.71 | 96.13 | 96.55 |
| Allowed (%) | 3.25 | 3.70 | 3.29 |
| Disallowed (%) | 0.03 | 0.16 | 0.16 |

## Notes

### Competing Interest Statement

The authors have declared no competing interest.

